# Inhibition of infection-induced vascular permeability modulates host leukocyte recruitment to *Mycobacterium marinum* granulomas in zebrafish

**DOI:** 10.1101/2021.11.23.469781

**Authors:** Julia Y Kam, Tina Cheng, Danielle C Garland, Warwick J Britton, David M Tobin, Stefan H Oehlers

## Abstract

Mycobacterial granuloma formation involves significant stromal remodeling and the growth of leaky, granuloma-associated vasculature. These permeable blood vessels aid mycobacterial growth, as anti-angiogenic or vascular normalizing therapies are beneficial host-directed therapies in pre-clinical models of tuberculosis. Here we demonstrate that vascular normalization through vascular endothelial-protein tyrosine phosphatase inhibition decreases granuloma hypoxia, the opposite effect of hypoxia-inducing anti-angiogenic therapy. Vascular normalization leads to increased T cell and decreased neutrophil recruitment to granulomas, correlates of a protective immune response against mycobacterial infection.

## Background

Mycobacteria induce changes in host vasculature, including angiogenesis and vascular permeability, through the production of a modified form of trehalose dimycolate [1–8]. Trehalose dimycolate triggers the expression of vascular endothelial growth factor (VEGF) by host cells, including macrophages [8]. The VEGF ligand signals through endothelial cell vascular endothelial growth factor receptor (VEGFR) to stimulate angiogenesis and increase vascular permeability [8–10]. Anti-angiogenic host-directed therapies control mycobacterial growth by preventing the delivery of oxygen to granulomas and increasing granuloma hypoxia [2, 10]. The Angiopoietin/Tie2 signaling pathway is a second vasoactive pathway that controls mycobacterial infection-induced vascular permeability [1]. Expression of the Angiopoietin 2 (ANG-2) ligand is upregulated in mycobacterial infection, allowing it to bind to and activate the TIE2 receptor. Increased ANG-2/TIE2 signaling antagonizes the stabilizing effects of ANG-1/TIE2 binding through Vascular endothelial protein tyrosine phosphatase (VE-PTP)-mediated deactivation of TIE2, resulting in increased vascular permeability [11–13]. Pharmacological inhibition of VE-PTP to reduce infection-induced vascular permeability results in reduced mycobacterial infection in the zebrafish model [1]. However, the mechanism connecting vascular integrity and the outcome of mycobacterial infection is unknown.

Experiments in murine cancer models have shown that normalization of blood vessel structure and function within solid tumors is associated with improved anti-tumor immunity [14–16]. Vascular normalization increases the ratio of T cell to neutrophil infiltration into tumors, which is indicative of an improved prognosis and results in favorable responses to immunotherapy. Due to the morphological similarities of mycobacterial granulomas and tumors, most importantly their tendency to develop hypoxia, we hypothesize that a similar mechanism of action may explain the protective effect of vascular normalization during mycobacterial infection [1–3, 9, 10].

The zebrafish-*M. marinum* infection system is an ideal model to investigate the relationship between changes to the granuloma stroma and the host immune response; granulomas reminiscent of *human-Mycobacterium tuberculosis* granulomas readily form in a system that is both genetically and pharmacologically tractable for *in vivo* studies [17]. The adult zebrafish allows the study of innate and adaptive immune cells, while the live imaging capability of zebrafish embryos and larvae allows visualization of innate immune cell behavior in intact animals. The zebrafish has been particularly important in the study of granuloma-associated vascular pathologies that impact the outcome of infection including angiogenesis, vascular permeability, and hemostasis [1, 3, 10, 18].

In this study, the zebrafish-*M. marinum* infection model was used to interrogate the effect of vascular normalization on immunity to mycobacterial infections using the previously characterized VE-PTP inhibitor AKB-9785 to normalize the vasculature.

## Methods

### Zebrafish handling

Adult zebrafish were held at the Garvan Institute of Medical Research Biological Testing Facility (St Vincent’s Hospital AEC Approval 1511), and the Centenary Institute (Sydney Local Health District AWC Approval 17-036). Experiments on adult zebrafish were carried out in accordance with Sydney Local Health District AWC Approval 16-037.

Transgenic strains used were: *TgBAC(lck:gfp)^vcc6^* [19], *Tg(lyzC:dsred)^nz50^* and *Tg(lyzC:gfp)^nz117^* [20], *Tg(kdrl:egfp)^s843^* [21].

Embryos were reared in E3 media at 28°C and supplemented with phenylthiourea (Sigma-Aldrich) to inhibit pigmentation from 1 day post fertilization (dpf).

### Experimental infection of adults

Adult zebrafish aged older than 3 months post fertilization were infected with ~200 CFU *M. marinum* M strain by intraperitoneal injection as previously described [22]. Briefly, adult fish were anaesthetized in tricaine and a 15 μl volume of bacterial inoculum was injected into the anaesthetized adult fish via intraperitoneal injection with a 28 G needle and syringe. Injected fish were recovered in 1 g/L salt water or aquarium system water. Infected fish were housed in a 28 °C incubator with 14 hour light:10 hour dark cycle, and were fed and monitored daily. Adult infected fish were euthanized by tricaine overdose at timepoints indicated for the collection of histology specimens.

### Drug Treatment

Adult zebrafish were immersed in 50 μM AKB-9785 (Aerpio Therapeutics) dissolved in dimethyl sulfoxide (DMSO) or the equivalent volume of DMSO vehicle control at time points indicated to a final concentration of 0.1% DMSO. Embryos were immersed in 12.5 μM AKB-9785 to a final concentration of 0.025% DMSO. Drug was refreshed every second day for adults and embryos.

### Hypoxyprobe staining

Hypoxyprobe staining was carried out as previously described [10, 22]. Briefly, adult zebrafish were injected with 15 μl of a 10 mg/ml pimonidazole solution (HP7; Hypoxyprobe) every two days from 7 days post infection (dpi) to 13 dpi and then euthanized for histology at 14 dpi.

### Histology

Euthanized adult fish were fixed in 10% neutral buffered formaldehyde (NBF) for 1-2 days at 4 °C. Fixed specimens were washed in PBS and processed for cryosectioning in OCT as previously described [22]. Frozen specimens were sectioned at 20 μm with a Leica Cryostat.

Hypoxyprobe was detected by staining with 4.3.1.3 mouse Dylight 549-MAb (HP7; Hypoxyprobe) or with unconjugated 4.3.1.3 mouse MAb and secondary detection with goat anti-mouse Alexa-Fluor 647 (A-21235; Life Technologies). Immunofluorescence was used to boost GFP signal in *TgBAC(lck:gfp)^vcc6^* specimens with a chicken anti-GFP primary (Abcam: ab13970) and goat anti-Chicken IgY (H+L), Alexa Fluor^®^ 488 secondary (Abcam: ab150169).

### Imaging

Histological sections of adult zebrafish were counterstained with DAPI and imaged with a Leica DM6000B. Static live imaging of zebrafish embryos was carried out on a Leica M205FA fluorescent stereo microscope. Time lapse imaging of zebrafish embryos was carried out using a Deltavision Elite High-resolution microscope.

Image analysis was carried out by fluorescent pixel count using ImageJ where appropriate [22, 23]. Tracking of neutrophil extravasation and migration was carried out manually in ImageJ.

### Infection of embryos

Microinjection was carried out as previously described [1, 10, 24]. Briefly, approximately 200 CFU of *M. marinum* M strain was microinjected into the neural tube or “trunk” of 2 dpf embryos.

### Tail wound neutrophil recruitment assay

Zebrafish embryos were anaesthetized, and tails were amputated using a surgical scalpel to induce a sterile wound. Embryos were immediately recovered into media containing AKB-9785 or DMSO and imaged at 6 hours post wounding.

### Statistical analysis

Statistical analysis was performed using GraphPad Prism 9. Statistical tests were carried out as indicated, the error bars represent standard deviation and P-values are supplied in the figures.

## Results

### Treatment with AKB-9785 reduces *M. marinum* burden as measured by histology

We first sought to validate our previously published report that AKB-9785 treatment of an existing *M. marinum* infection reduced the culturable bacterial burden using histological methods [1]. Analysis of total fluorescent pixel counts from census cryosections of adult zebrafish revealed reduced *M. marinum* burden (Figure 1A).

**Figure 1:**
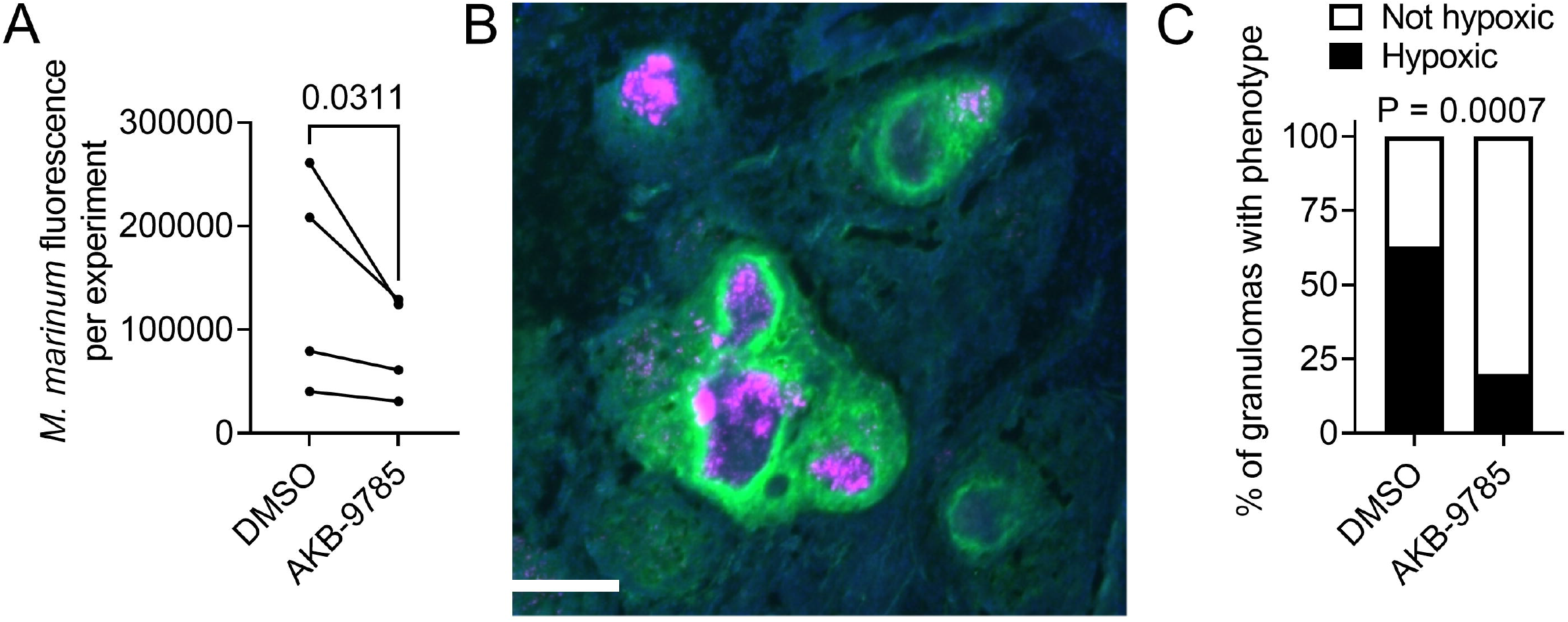
Vascular normalization reduces *M. marinum* granuloma hypoxia in adult zebrafish. A. Quantification of average bacterial burden by histological fluorescent pixel count across 4 paired experiments with at least 3 animals. Statistical testing by ratio paired T test. B. Representative image of a cluster of hypoxyprobe-positive necrotic granulomas from a 2 wpi *M. marinum*-infected zebrafish. Scale bar represents 100 μm. C. Quantification of granuloma hypoxia by hypoxyprobe staining. Total granulomas analyzed DMSO = 27 and AKB-9785 = 35; total animals analyzed: DMSO = 3 and AKB-9785 = 3. Statistical testing by Fisher’s exact test on raw counts.

### Treatment with AKB-9785 reduces granuloma hypoxia

Vascular normalization by VEGF blockade has been demonstrated to reduce *M. tuberculosis* granuloma hypoxia in rabbits [4]. We performed hypoxyprobe staining of adult zebrafish that had been soaked in DMSO and AKB-9785 from 1 to 2 wpi to quantify the proportion of hypoxic granulomas in our system (Figure 1B). Treatment with AKB-9785 reduced the proportion of hypoxic granulomas (Figure 1C), confirming the fidelity of our model.

### Vascular permeability correlates with neutrophil recruitment to *M. marinum* granulomas in adult zebrafish

As vascular normalization by modulation of endothelial junctional permeability had been reported to reduce the recruitment of neutrophils to solid tumors and neutrophil influx has been correlated with progressive histopathology in human TB [14–16], we hypothesized that treatment with AKB-9785 would also reduce neutrophil recruitment to *M. marinum* granulomas in zebrafish.

Adult *Tg(lyzC:dsred)^nz50^* zebrafish, where neutrophils are labelled by DsRed expression [25], were infected with *M. marinum* were treated with AKB-9785 from 1 to 2 wpi and cryosectioned to quantify the response of neutrophil recruitment to vascular normalization (Figure 2A). Treatment with AKB-9785 increased the area of bacterial fluorescence per granuloma by a small but statistically significant margin (Figure 2B), decreased neutrophil recruitment to granulomas by approximately 30% (Figure 2C), resulting in a reduced neutrophil to bacteria ratio per granuloma (Figure 2D).

**Figure 2:**
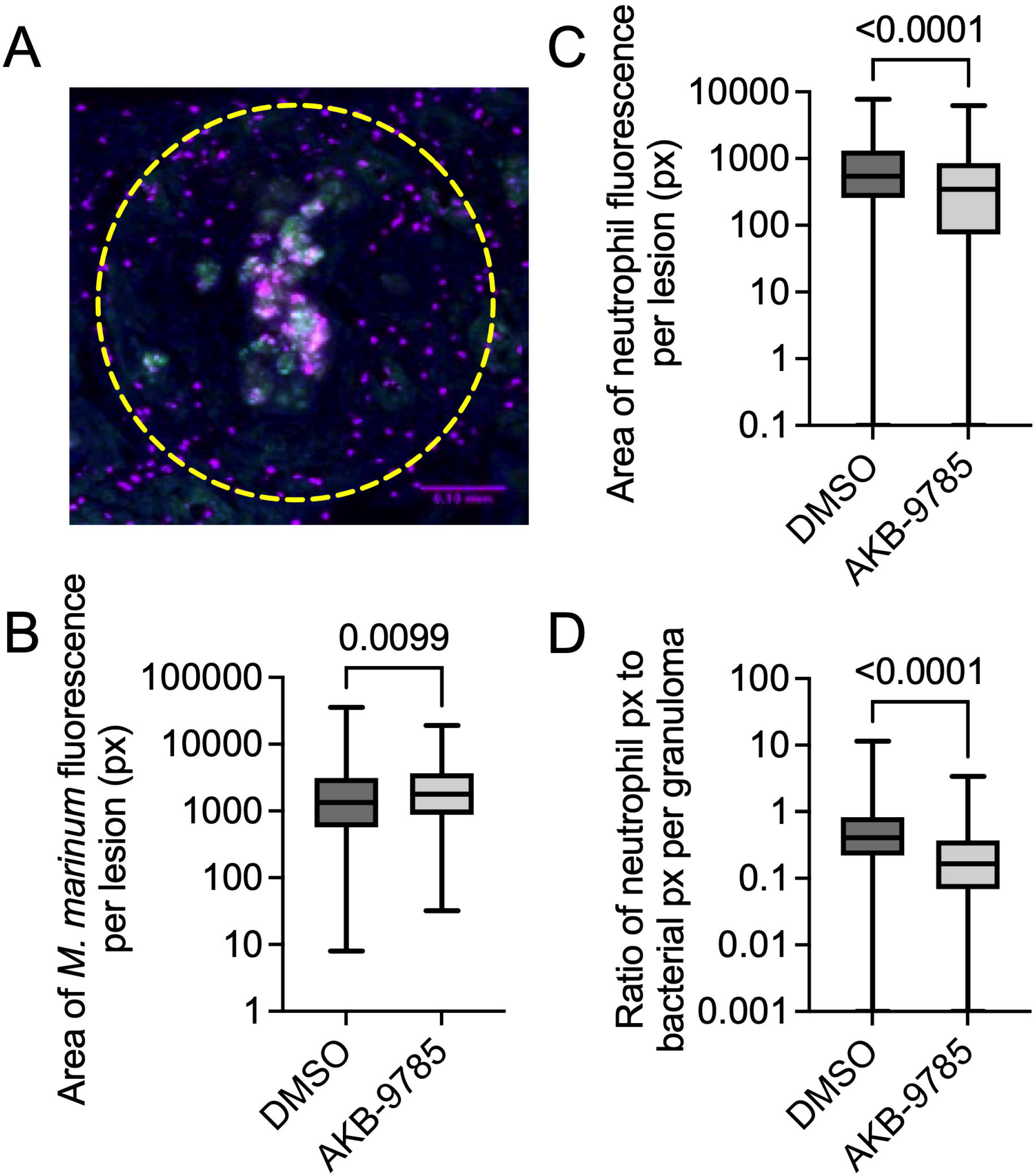
Vascular normalization reduces neutrophil recruitment to *M. marinum* infection in adult zebrafish. A. Representative image of a DAPI-stained (blue) necrotic granuloma from a 2 wpi *Tg(lyzC:dsred)^nz50^* (magenta) adult zebrafish infected with *M. marinum*-wasabi (green). Yellow dashed region indicates area measured for quantification. Scale bar represents 100 μm. B. Quantification of granuloma bacterial load by fluorescent pixel count. Total granulomas analyzed DMSO = 324 and AKB-9785 = 187; total animals analyzed: DMSO = 4 and AKB-9785 = 4. C. Quantification of neutrophil recruitment to granulomas by fluorescent pixel count. D. Calculation of neutrophil to bacterial fluorescent pixel count ratio. Statistical tests for B, C, D were performed using Mann-Whitney U-test. Data are representative of two experimental replicates with similar numbers of animals.

### Vascular normalization reduces neutrophil recruitment to *M. marinum* granulomas in zebrafish embryos

To further investigate the relationship between AKB-9785 treatment-induced vascular normalization and neutrophil recruitment we sought to image neutrophil extravasation live in optically transparent zebrafish embryos at 7 dpi (Figure 3A). Treatment with AKB-9785 did not significantly affect the overall bacterial burden or neutrophil recruitment across all analyzed granulomas (Figure 3B and 3C). However, AKB-9785 treatment did significantly reduce the neutrophil to bacteria ratio per granuloma, suggesting that vascular normalization reduced neutrophil recruitment to granulomas in zebrafish embryos (Figure 3D).

**Figure 3:**
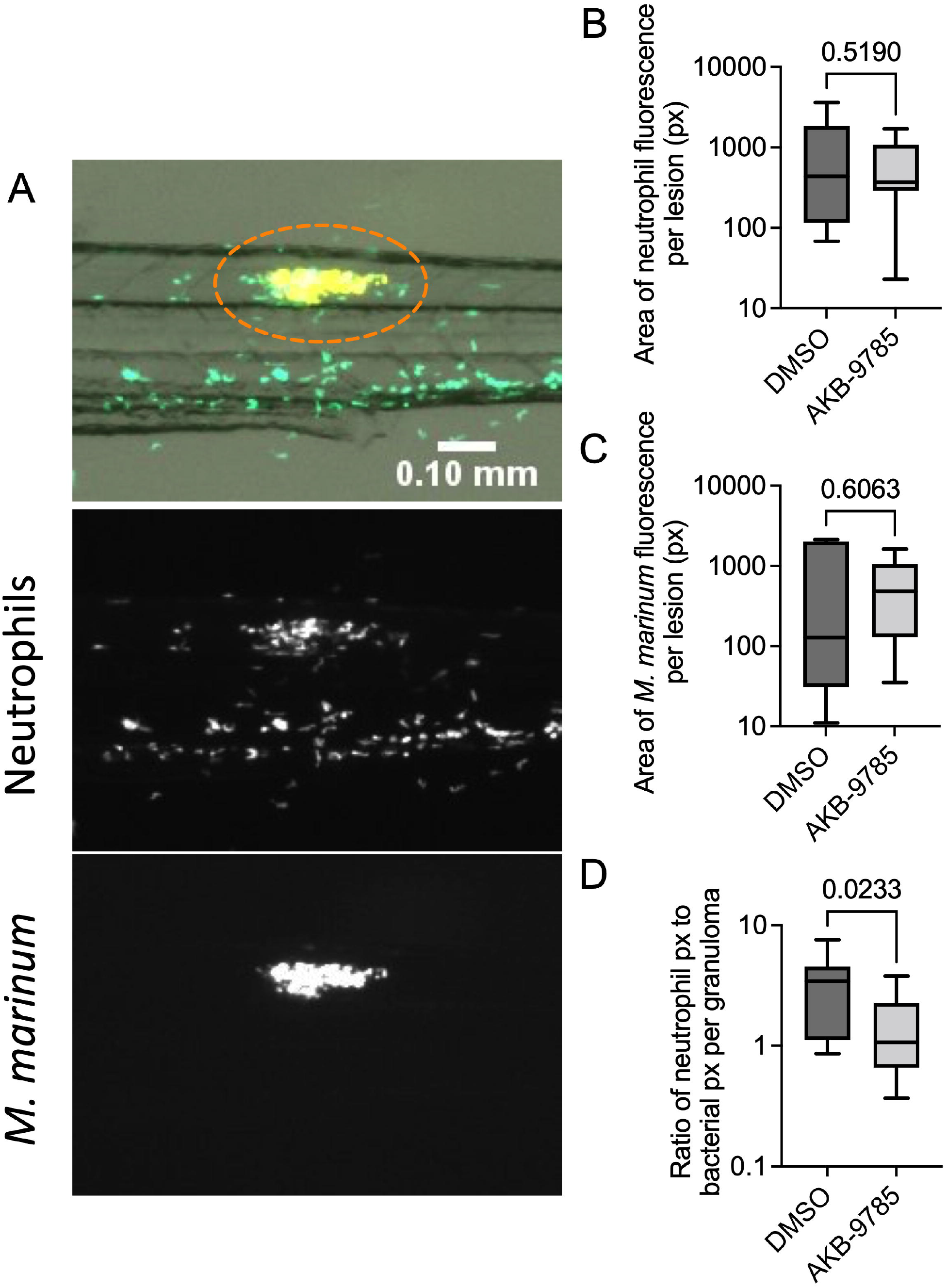
Vascular normalization reduces neutrophil recruitment to *M. marinum* infection in zebrafish embryos. A. Representative image of 7 dpi *M. marinum*-TdTomato (appears yellow due in overlay) infected *Tg(lyzC:gfp)^nz117^* zebrafish embryo with pseudo coloring of neutrophils green. Orange dotted lines in overlay image outline the area quantified. Scale bar represents 100 μm. B. Quantification of granuloma bacterial load by fluorescent pixel count. Total granulomas analyzed DMSO = 12 and AKB-9785 = 13; total animals analyzed: DMSO = 5 and AKB-9785 = 5. C. Quantification of neutrophil recruitment to granulomas by fluorescent pixel count. D. Calculation of neutrophil to bacterial fluorescent pixel count ratio. Statistical tests for B, C, D were performed using Mann-Whitney U-test. Data are representative of two experimental replicates with similar numbers of animals.

### Vascular normalization focuses the local extravasation of neutrophils

We next used live imaging to investigate if vascular normalization affected the extravasation of granuloma-associated neutrophils (Figure 3A). Neutrophil extravasation was scored as either proximal, occurring within two intersegmental vessels of the granuloma, or distal, occurring more than two intersegmental vessels away from the granuloma. Treatment with AKB-9785 increased the proportion of recruited neutrophils from proximal blood vessels (Figure 4B). Interestingly, AKB-9785 treatment reduced the point-to-point velocity of neutrophils that extravasated from proximal blood vessels, but no change was seen in neutrophils that extravasated from distal blood vessels (Figure 4C).

**Figure 4:**
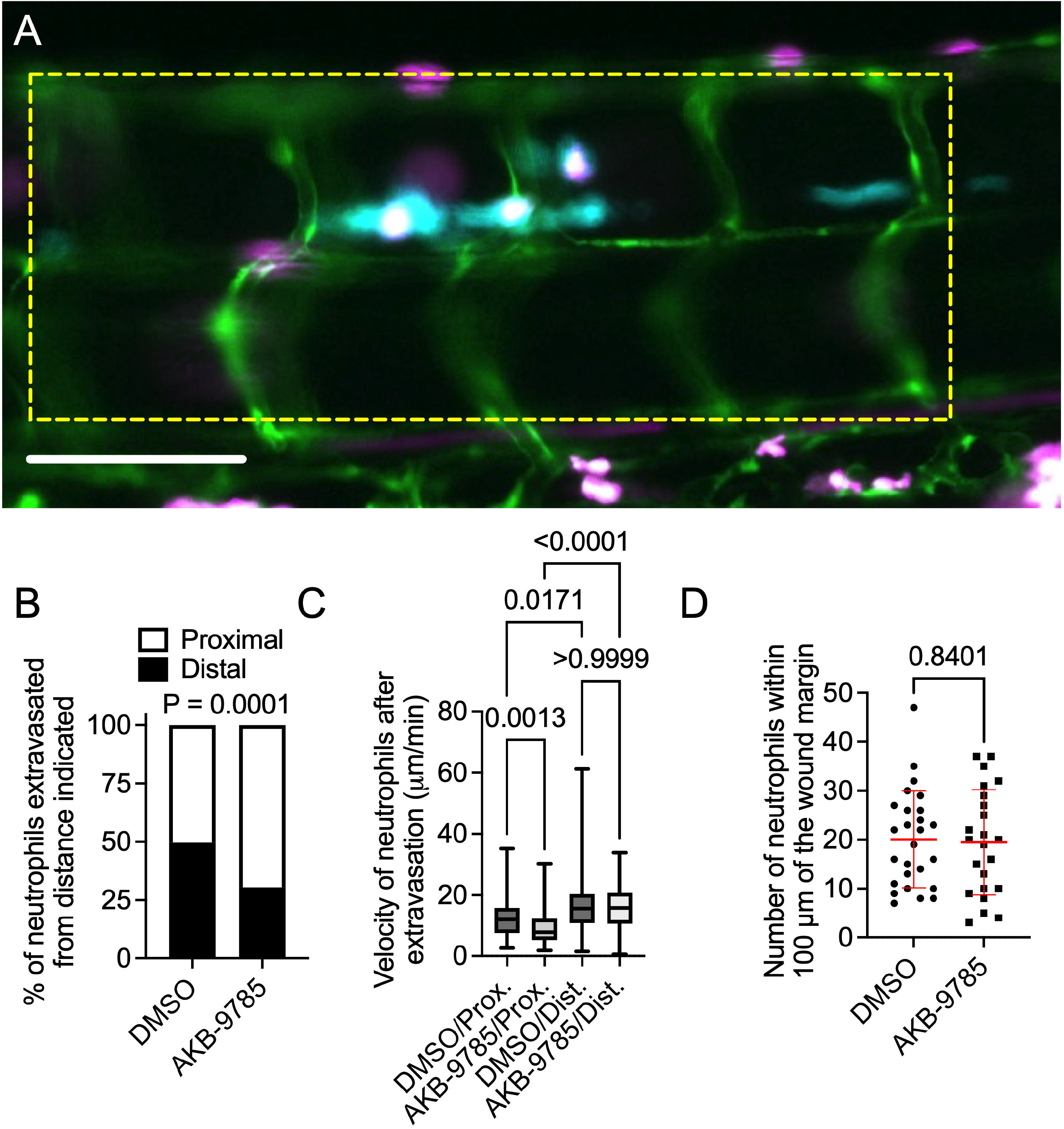
Vascular normalization focuses neutrophil extravasation to blood vessels near granulomas in zebrafish embryos. A. Representative image of a granuloma from a 4 dpi *M. marinum*-Cerulean (blue) infected *Tg(Tg(kdrl:egfp, lyzC:dsred)^s843, nz50^* zebrafish embryo with pseudo coloring of blood vessels green and neutrophils magenta. Yellow dotted box outlines the two intersegmental vessel bounds around the central granulomas used to define proximal extravasation events. Scale bar represents 100 μm B. Quantification of neutrophil extravasation event source for neutrophils recruited to *M. marinum* granulomas, proximal defined as within 2 intersegmental vessels. Total number of events measured: DMSO = 153, AKB-9785 = 262; total embryos imaged: DMSO = 8, AKB-9785 = 12. C. Quantification of neutrophil straight-line velocity after extravasation until neutrophil reaches *M. marinum* granuloma. D. Counts of neutrophils within 100 μm of the tail amputation margin at 6 hours post wounding. Each data point represents a single animal. Statistical test for B was performed by Fisher’s exact test on raw counts, statistical test for C was performed by Mann-Whitney U-test, statistical test for D was performed using unpaired T-test. D data are representative of two experimental replicates with similar numbers of animals.

To investigate if treatment with AKB-9785 had a direct effect on neutrophil recruitment we performed tail wound experiment to quantify neutrophil recruitment in a system with direct interstitial neutrophil recruitment. No significant differences were observed between DMSO and AKB-9785 soaked animals (Figure 4D), suggesting that AKB-9785 acts to limit neutrophil recruitment only in the context of infection-induced vascular permeability.

### Vascular normalization increases T cell recruitment to *M. marinum* granulomas

The adult zebrafish contains an adaptive immune system with many similarities to that of mammals, making it an ideal animal model for the study of the adaptive immune response to natural mycobacterial infection. To study the effect of vascular normalization on T cell infiltration of *M. marinum* granulomas, we infected *TgBAC(lck:gfp)^vcc4^* zebrafish where T cells are labeled with GFP expression with *M. marinum-TdTomato* and treated with AKB-9785 from 1 to 2 wpi followed up cryosectioning to quantify the response of T cells to vascular normalization (Figure 5A).

**Figure 5:**
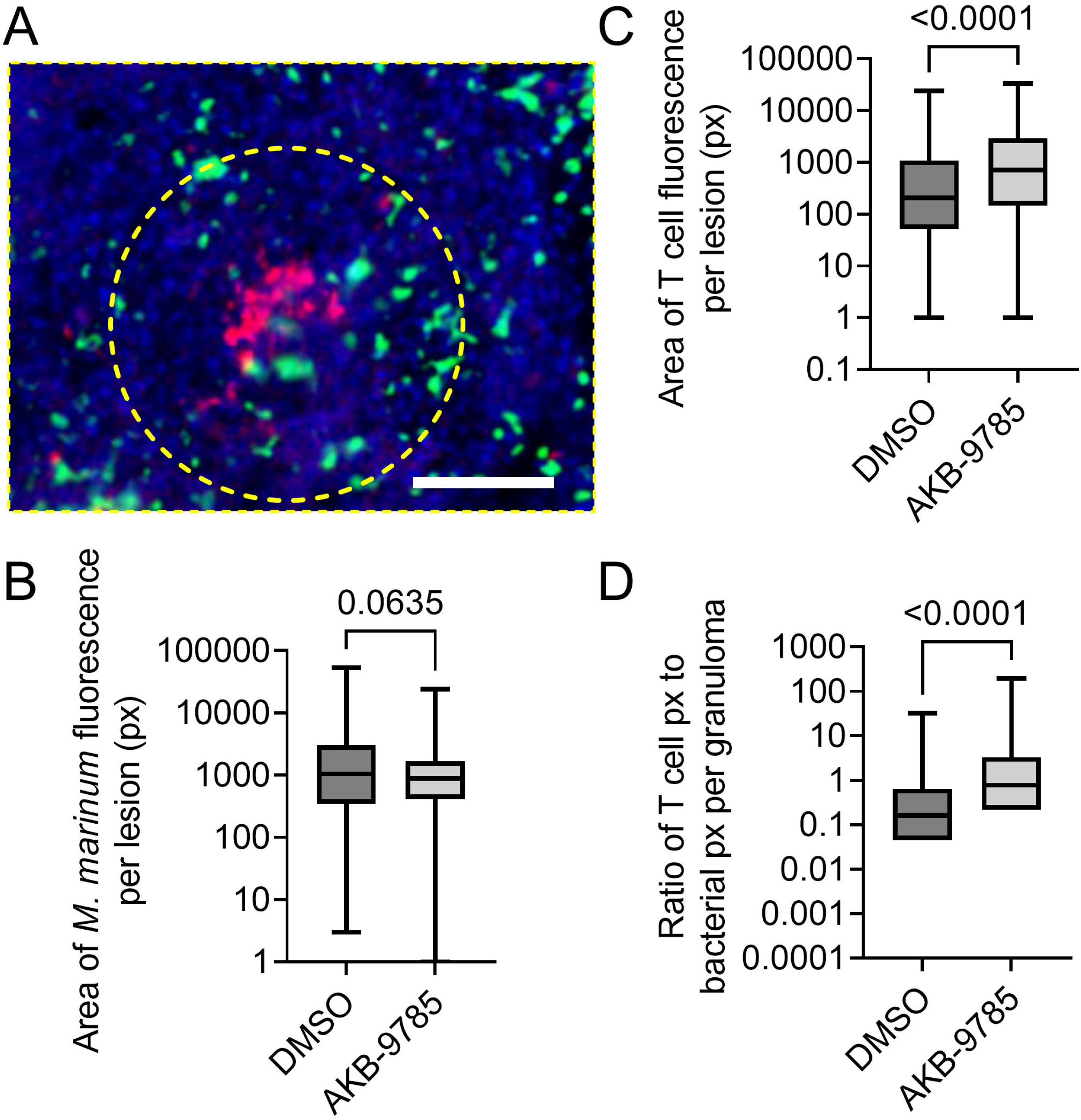
Vascular normalization increases T cell recruitment to *M. marinum* infection in adult zebrafish. A. Representative image of a cellular granuloma from a 2 wpi *TgBAC(lck:gfp)^vcc6^* adult zebrafish infected with *M. marinum*-Katushka. Yellow dashed region indicates area measured for quantification. Scale bar represents 100 μm. B. Quantification of granuloma bacterial load by fluorescent pixel count. Total granulomas analyzed DMSO = 324 and AKB-9785 = 187; total animals analyzed: DMSO = 3 and AKB-9785 = 3. C. Quantification of T cell recruitment to granulomas by fluorescent pixel count. D. Calculation of T cell to bacterial fluorescent pixel count ratio. Statistical tests for B, C, D were performed using Mann-Whitney U-test. Data are representative of three experimental replicates with similar numbers of animals.

Treatment with AKB-9785 did not affect the area of bacterial fluorescence per granuloma (Figure 5B), but increased T cell recruitment to granulomas (Figure 5C), resulting in an increased T cell to bacteria ratio per granuloma (Figure 5D).

## Discussion

Our studies demonstrate that vascular permeability alters neutrophil and T cell recruitment to mycobacterial granulomas, suggesting that infection-induced vascular permeability may facilitate unfavorable neutrophilic inflammation while preventing effective T cell recruitment. Furthermore, our observation that vascular normalization reduced granuloma hypoxia in our model fits with existing literature that neutrophils are retained in hypoxic lesions while T cells are repelled [2, 26–28].

The importance of the zebrafish adaptive immune system has been previously established using *rag* knockout animals, which rapidly succumb to *M. marinum* infection [29]. Here, we have demonstrated the converse: that increasing T cell recruitment to *M. marinum* granulomas correlates with protection from infection. Our findings suggest that there is suboptimal T cell recruitment in natural *M. marinum* infection of zebrafish due to infection-induced vascular permeability and the induction of granuloma hypoxia.

While intensive infiltration of neutrophils is a negative prognostic marker in chronic TB, there is evidence from zebrafish models that neutrophils are beneficial early and when granuloma macrophage epithelization is blocked [26, 30]. Thus, it is not clear why reducing the number of neutrophils recruited to granulomas is beneficial in the embryo which lacks T cells to explain the responsiveness to vascular normalization therapy. While it is possible that the effect seen in embryos is due to another aspect of vascular normalization, we hypothesize that vascular normalization minimizes the interstitial tissue “commuting” distance of extravasated neutrophils by reducing the distal extravasation of neutrophils, thereby reducing the amount of neutrophil-generated immunopathology caused by distally extravasated neutrophils migrating to granulomas [31, 32].

In addition to its effect on cellular immune responses, abnormal and dysfunctional vasculature in granuloma and tumor environments can also impair the delivery of antibiotics and this can greatly impair the effectiveness of antibiotic or anti-cancer therapy [9]. A study by Xu et al. demonstrated that inhibition of matrix metalloproteinase (MMP) increased pericyte coverage in blood vessels, stabilizing the integrity of *M. tuberculosis* infected mouse lung tissue, which in turn enhanced the *in vivo* potency of frontline tuberculosis drugs such as isoniazid and rifampicin [33]. Vascular normalization is thus more favorable as an adjunctive therapy for antibiotic-sensitive infection than anti-angiogenic therapies, which may increase the exclusion of antibiotics from granulomas.

Neutrophil tail wound and migration velocity after extravasation from distal blood vessels were the same after AKB-9785 treatment, suggesting AKB-9785 does not act directly on neutrophil motility to limit their infiltration into *M. marinum* granulomas. We did observe a decrease in the velocity of proximally extravasated neutrophils in AKB-9785 treated embryos consistent with the possibility that the embryonic granulomas were less hypoxic and thus less “attractive” to neutrophils. While expression of the AKB-9785 target VE-PTP is considered endothelial-specific, the report that VEGF directly modulates the recruitment of immune cells in mouse mycobacterial infection models leaves the possibility that AKB-9785 may indirectly modulate immune cell recruitment through downstream cytokines [7].

Overall, this study provides evidence for the use of vascular normalization to enhance a more effective immune response in mycobacterial infections. An increased ratio of T-cell to neutrophil infiltration is frequently a favorable prognostic marker both in human infections, such as tuberculosis, and in cancer. Neutrophilic inflammation and lack of T cell recruitment is a particularly important component of the immunopathology in respiratory viral infections, such as influenza [34]. We hypothesize that the use of vascular normalization strategies including VEGF or VEGFR blockade or inhibition [16], targeted increase in VE-cadherin expression [14], dual ANG2-blocking and TIE2-activating antibodies and the VE-PTP inhibition used in this study will be of benefit to a wide range of infectious diseases [35].

## Acknowledgements

The authors acknowledge members of the Tuberculosis Research Program at the Centenary Institute for discussion of the manuscript, Dr Angela Fountaine of the BioImaging Facility and Sydney Cytometry at Centenary Institute for technical assistance with imaging, and Aerpio Therapeutics for AKB-9785.

